# Improving RT-LAMP Detection of SARS-CoV-2 RNA through Primer Set Selection and Combination

**DOI:** 10.1101/2021.07.23.453545

**Authors:** Yinhua Zhang, Nathan A. Tanner

## Abstract

Reverse transcription loop-mediated isothermal amplification (RT-LAMP) has emerged as a viable molecular diagnostic method to expand the breadth and reach of nucleic acid testing, particularly for SARS-CoV-2 detection and surveillance. While rapidly growing in prominence, RT-LAMP remains a relatively new method compared to the standard RT-qPCR, and contribution to our body of knowledge on designing LAMP primer sets and assays can have significant impact on its utility and adoption. Here we evaluate 18 LAMP primer sets for SARS-CoV-2, comparing speed and sensitivity with different LAMP formulations and conditions across more than 5,000 RT-LAMP reactions and identifying several primer sets with similar high sensitivity for different SARS-CoV-2 gene targets. Significantly we observe a consistent sensitivity enhancement by combining primer sets for different targets, confirming and building on earlier work to create a simple, general approach to building better and more sensitive RT-LAMP assays.

## Introduction

The ongoing COVID-19 pandemic has brought an urgent demand for molecular diagnostic testing at an unprecedented scale. Reverse transcription quantitative PCR (RT-qPCR) has long been the standard for molecular testing and has been widely and massively used for SARS-CoV-2 detection. However, due to the large volume of testing required worldwide (e.g. 1-2 million daily tests in the US alone) and the need for testing outside of clinical laboratories, molecular diagnostic methods other than RT-qPCR have also become widely used. Isothermal amplification methods such as transcription mediated amplification (TMA) and nicking enzyme amplification reaction (NEAR) are the core chemistry for test platforms from Hologic and Abbott, respectively, with other methods showing promise for potential widespread testing. Of these alternatives, reverse transcription loop-mediated isothermal amplification (RT-LAMP) has now been used most prominently [1, 2] with numerous diagnostic tests based on RT-LAMP receiving Emergency Use Authorization from the FDA including the first ever at-home, over-the-counter molecular diagnostic test from Lucira Health [3–6].

The widespread interest in RT-LAMP for molecular detection of SARS-CoV-2 has generated a lot of interest and information on how to achieve efficient and reliable detection. LAMP is a much newer method than PCR, with protocols and assay design methods not quite as established, though the recent increase in LAMP development efforts is rapidly changing this discrepancy. Many factors can have an impact on LAMP assay performance: sample source (nasopharyngeal or nasal swab, saliva); sample processing (direct sample or purified RNA); and uniquely for LAMP the detection modality and instrument/device design, e.g. pH-based colorimetric, fluorescence, Cas enzyme cleavage, etc. And regardless of test design, a critical factor to performance is of the assay primers and reaction conditions for sensitive RNA detection.

Typical RT-LAMP reactions use a primer set covering 8 regions on the target sequence and the primer design is facilitated by software (Primer Explorer V5 at Eiken https://primerexplorer.jp/e/, or LAMP Primer Design Tool at NEB https://lamp.neb.com/#!/) based on oligo length, AT and GC content and thermodynamic stability. However, like primers for RT-qPCR, not all software-designed primers perform optimally and it is often necessary to screen several sets to obtain ones that give satisfactory specificity and sensitivity. Among the published works evaluating RT-LAMP for SARS-CoV-2 detection, most screened many sets of primers before deciding on one set to continue their study. For example, 35 sets were screened by Yang et al [7] to obtain 3 sets targeting 3 different genomic regions for conventional RT-LAMP reaction; 29 sets were screened by Joung et al [6] to a single set that worked well in a coupled one-pot RT-LAMP/CRISPR cleavage assay.

To date, many sets of SARS-CoV-2 LAMP primers have been published and some used in EUA or CE-IVD diagnostic tests. Comparison of these assays and primer sets based on published data can be challenging, as methods, reagents, template sources, and other differences may have effects on sensitivity and complicate data interpretation. Here we describe evaluating RT-LAMP primer sets under identical conditions for a fairer comparison of that critical reaction parameter. Similar analyses performed previously have suggested a wide range of sensitivity among published primer sets [8, 9]. Building on these studies we apply here a more stringent approach, performing large numbers of repeats and evaluating performance with different RT-LAMP reagents and amplification temperatures, running >5,000 RT-LAMP reactions to more fully evaluate the performance of SARS-CoV-2 assays. Significantly, we demonstrate a general principle to further increase LAMP detection sensitivity by combining primer sets in the same reactions, expanding on our previous observations [10].

## Materials and Methods

### RT-LAMP primer selection

17 LAMP primer sets from previous publications and 1 new primer set were selected for evaluation (Table 1). 3 sets (N2, E1 and Ase1) were studied in our previous publication [10]. 10 sets (S2 [11], S4 [12], S10 [12], S11 [13], S12 [3], S13 [7], S14 [14], S17 [12], S18 [15], Mam-N/S19[3]) were shown to be the most sensitive ones in a previous analysis of 19 published sets [8]. 5 primer sets (S-Huang[16], N-Baek [17], As1e [18], S-Yan [13], N-Lu [15]) were screened as the most sensitive sets out of 16 published sets [9]. These included 2 sets (S-Yan=S11, N-Lu=S18) that overlapped with those selected from Dong et al. A new set (SGF-wt) was selected based on sensitive detection at a region of SARS-CoV-2 variant sequence deletion in our testing. Primer set Joung was from Joung et al [6] and Lau from Lau et al [19].

All primers were synthesized by IDT at 100 or 250 nMole scale with standard desalting. For larger-scale oligo synthesis HPLC purity of the FIP and BIP is generally recommended for enhanced RT-LAMP performance, but does not seem to affect small scale oligo synthesis significantly. Primers were dissolved and then mixed in ddH2O as 25x stocks of each set based on standard 1x final concentrations in LAMP: 0.2 μM F3, 0.2 μM B3, 1.6 μM FIP, 1.6 μM BIP, 0.4 μM Loop F, 0.4 μM Loop B.

### RT-LAMP reactions

RT-LAMP reactions were performed using either WarmStart^®^ Colorimetric LAMP 2X Master Mix (DNA & RNA) (M1800) or WarmStart^®^ LAMP Kit (DNA & RNA) (E1700) from New England Biolabs (NEB). 40 mM guanidine hydrochloride was included in all reactions to improve LAMP reaction speed and sensitivity [10]. The same vial of synthetic SARS-CoV-2 RNA from Twist Bioscience (Twist Synthetic SARS-CoV-2 RNA Control 2 (MN908947.3), SKU: 102024) was used for all reactions, with SARS-CoV-2 RNA diluted and aliquoted in nuclease-free water and 10 ng/μl Jurkat total RNA (Biochain). RT-LAMP reactions were performed in 25 μl volumes supplemented with 1 μM SYTO^®^-9 double-stranded DNA binding dye (Thermo Fisher S34854) in 96-well plates and incubated at 65 or 60 °C on a realtime qPCR machine (BioRad CFX96). Amplification signal was acquired every 15 seconds for 108 “cycles” (total incubation time was ~40 min). The time to reach the signal threshold (Tt) was determined from the real time fluorescence signal and positive was scored using a cutoff of 22.5 minutes for RT-LAMP at 65 °C and 30 minutes at 60 °C. For each primer set or combination test condition, a minimum of 24 repeats were performed, and the number of positives were used as an indicator for detection sensitivity. At least 8 repeats of no-template control reactions were performed for each primer set or combination to evaluate production of target-independent signal. A low RNA template amount was used to ensure only a percentage of positive reactions in the repeats, but not in all in order to compare relative performance. To ensure a fair comparison and consistent reagent performance we used the same lots of reagents for all tests and included 24 repeats of N2 primer set with 50 copies of SARS-CoV-2 RNA at each assay date and confirmed consistent reaction times and positive detection results.

## Results

### Procedure to identify sensitive primer sets

RT-LAMP reactions were tested with both M1800 (pH-based colorimetric) and standard E1700 LAMP mixes. Both formulations were tested at 65 °C and 60 °C to capture any temperature preference by primers. We first tested single primer sets using 50 copies of SARS-CoV-2 synthetic RNA, with further evaluation of the most sensitive primer sets in various combinations and with 25 and 12.5 copies of RNA templates. The sensitivity of RT-LAMP was evaluated based on the number of positives in 24 repeats of reactions containing 50 copies of SARS-CoV-2 RNA (Figure 1). Each primer set was classified as sensitive, medium or poor based on the number of positives in the repeats. With M1800 colorimetric LAMP at 65 °C (Figure 1A), 8 primer sets (S4, S10, S11, S12, S13, S18, N2 and E1) were identified as sensitive, 7 medium (S2, S14, S17, Joung, As1e, SGF-wt and Mam-N) and 3 poor (N-Baek, S-Huang and N-Lau). With M1800 at 60 °C (Figure 1B), 7 sets (S10, S11, S12, S13, S17, N2 and E1) were classified as sensitive. 6 of those 7 sets were consistent at the two temperatures, but S4 and S18 showed high sensitivity at 65 °C that was reduced at 60 °C, and one set, S17, showed increased sensitivity at 60 °C.

**Figure 1.**
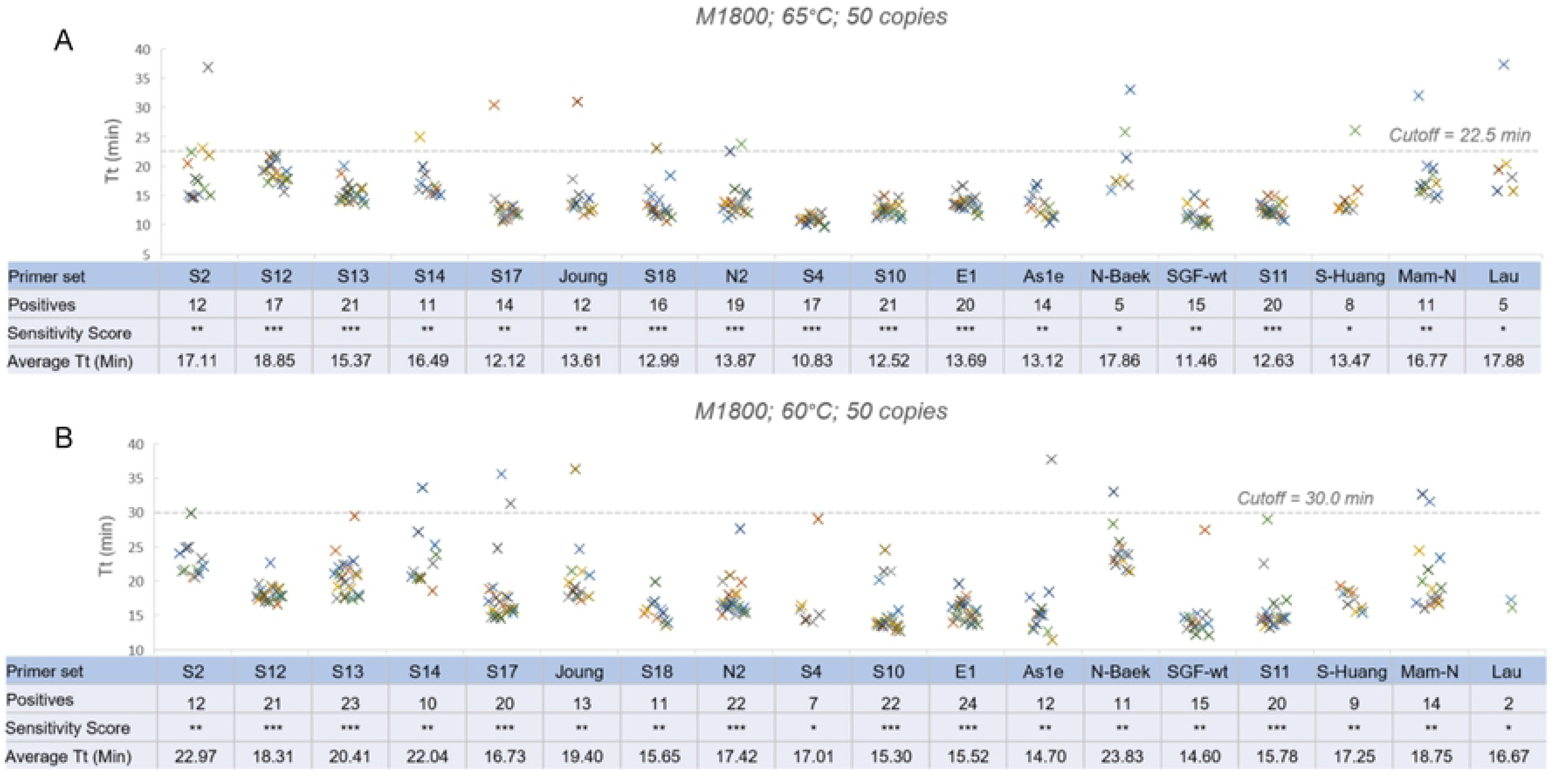
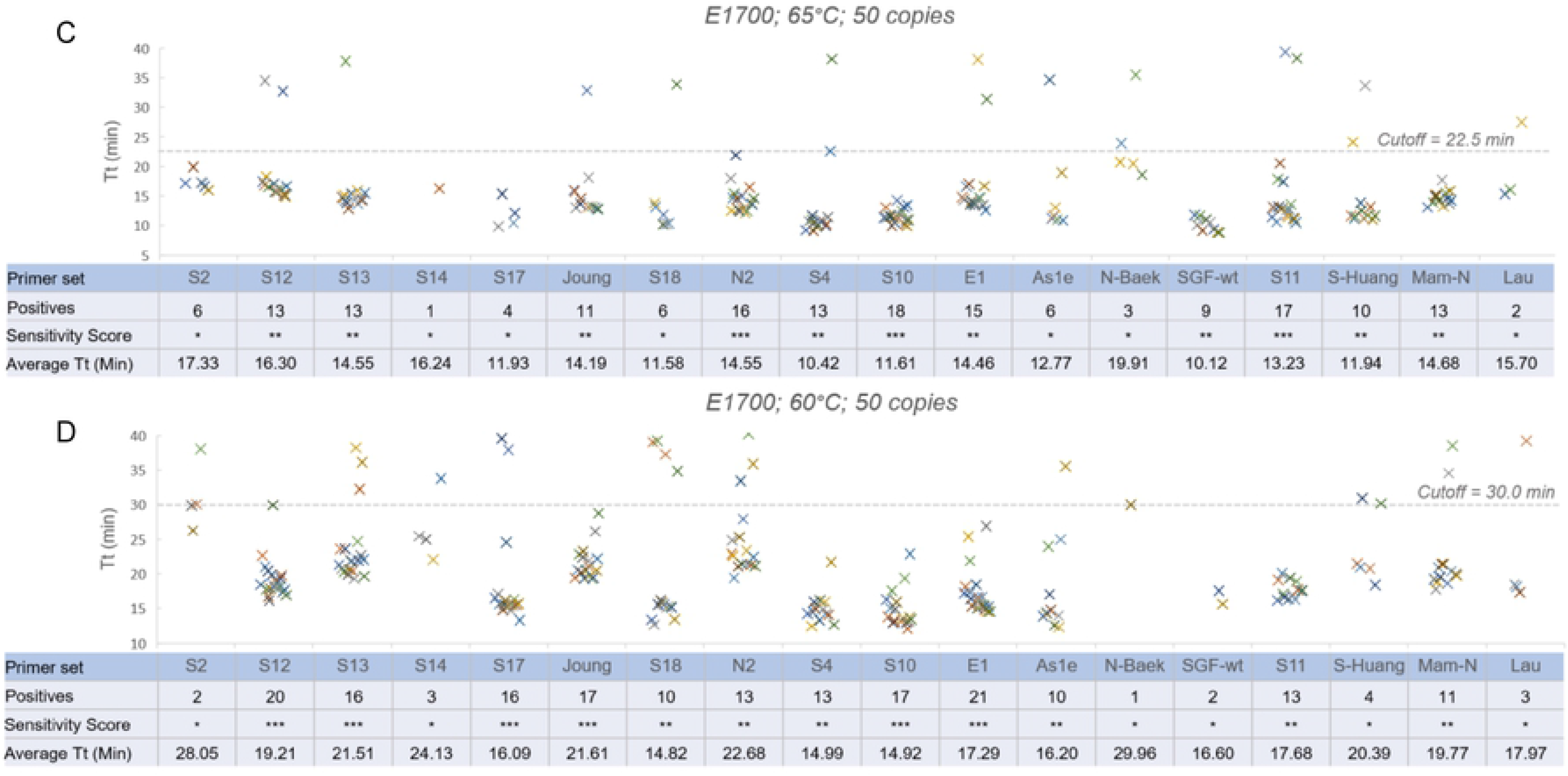
Performance of single primer sets. Each primer set was assayed with 24 repeats with 50 copies of SARS-CoV-2 RNA templates. Each primer set was assigned an sensitivity score: sensitive (*** 16-24 positives); medium (** 9-15 positives) or poor (* 1-8 positives). RT-LAMP reactions were conducted with M1800 Colorimetric LAMP at 65 °C (A) or 60 °C (B), and with E1700 LAMP Mix at 65 °C (C) or 60 °C (D).

RT-LAMP speed, indicated by the average Tt of positives, was generally pretty consistent across the 18 primer sets, ranging from 10.8–17.8 minutes with M1800 and 50 copies at 65 °C. There was some association with the faster primer sets and increased sensitivity, but correlation of positive detection with Tt was very weak (R^2^ = 0.23) implying that the two observables are not necessarily linked. Reaction speed for 16 of the 18 primer sets increased (by 3.9 ± 1.5 minutes) when temperature was reduced to 60 °C, with only S12 and Lau sets increasing slightly in speed; with E1700 all sets were slower at 60 °C. Taken together these results indicate that reaction speed is determined primarily by the polymerase amplification chemistry and not primer sequence.

When assayed with E1700, primer sets that gave high sensitivity with M1800 also showed higher sensitivity, however the overall positive numbers were slightly lower. There were 3 sets (S10, S11 and N2) in the sensitive category at 65 °C (Figure 1C) and 6 sets (S10, S12, S13, S17, E1 and Joung) in this group at 60 °C (Figure 1D). Primers that showed poor sensitivity in M1800 (N-Baek, S-Huang and N-Lau) as well as 5 additional sets (S2, S14, S17, S18 and As1e) perform poorly in E1700 at 65 °C. Among them, 3 sets (S17, S18 and As1e) showed improvement at 60 °C, moving to the medium category.

### Improving sensitivity by combining two sensitive primer sets

Sensitive primer sets were selected to test their performance in RT-LAMP reaction containing 2 primer sets. 5 sets (S10, S11, S12, S17, N2 and E1) were chosen based on high sensitivity using both RT-LAMP formulations and temperatures (S13 was one of the most sensitive sets but it was not selected here because its primers differ in only a few bases from primer set E1). RT-LAMP containing 2 primer sets was compared to a single primer set in the presence of 25 copies of SARS-CoV-2 RNA using M1800 at 65 and 60 °C (Figure 2A-B). In all the combinations tested (S10+S11; S10+S12; S11+S12; S12+S17; and N2+E1), the combined primer reactions gave higher sensitivity than with single primer sets at both temperatures. All combinations showed more sensitive detection compared to individual sets with both RT-LAMP formulations and at both 65 and 60 °C (Figure 2C-D). We next extended this comparison with even lower SARS-CoV-2 RNA templates (~12.5 copies). 6 primer sets were divided into two groups (S10, S11, S12; and N2, E1, As1e) and the single primer set was compared with one pair from each group (S10+S11, N2+E1). As1e was picked since it was assayed in our previous study [10] even though its detection sensitivity belonged to the medium category above. As shown, combination of 2 primer sets S10+S11 (Figure 3A-B) or N2+E1 (Figure 3C-D) also showed more sensitive detection than any single primer set even at this low level of templates in either M1800 or E1700 at 65 or 60 °C.

**Figure 2.**
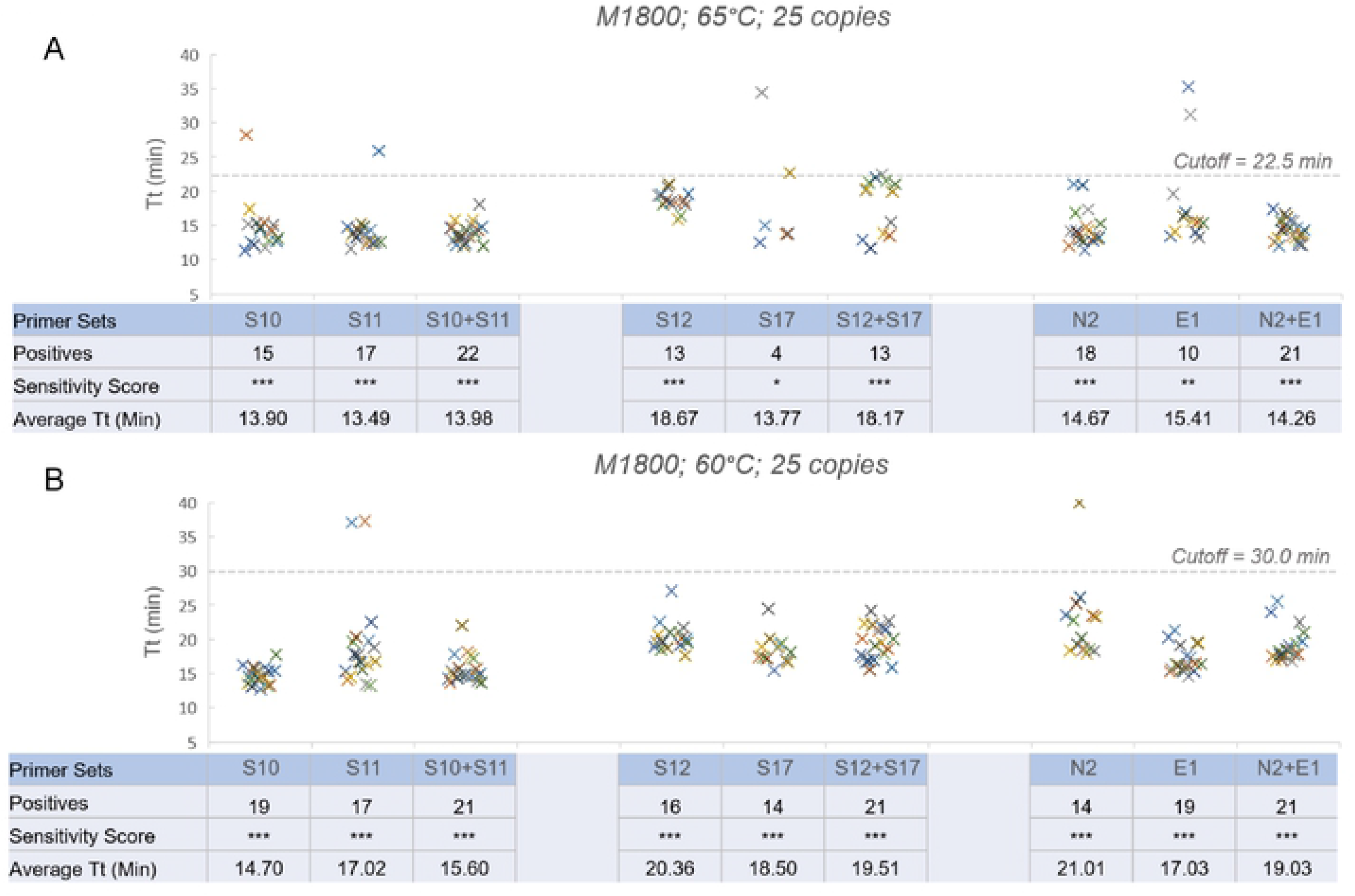
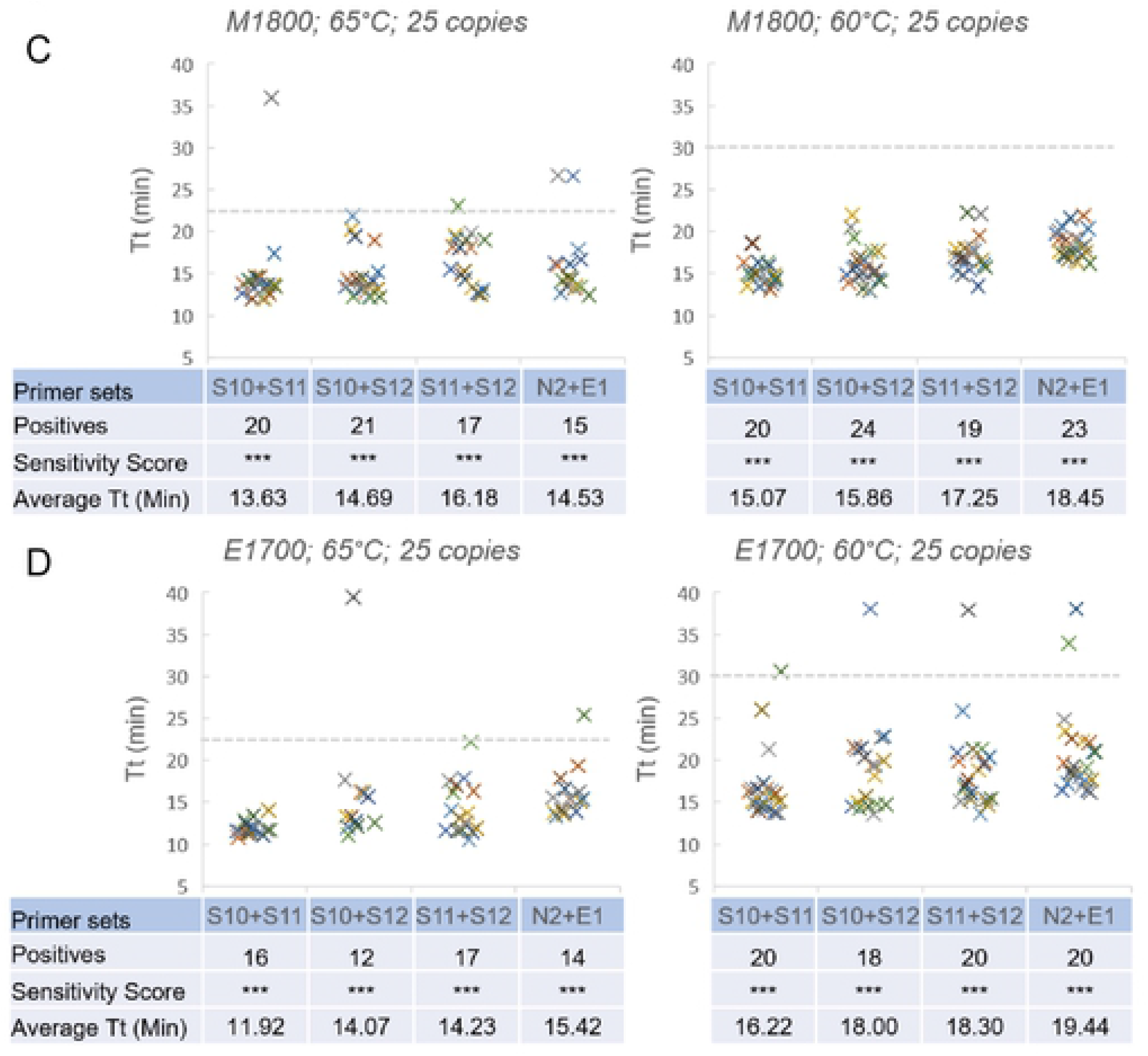
Increased detection sensitivity by combining 2 primer sets. Reactions with single primer sets were compared with that by a combination of 2 primer sets in the presence of 25 copies of SARS-CoV-2 RNA templates. The sensitivity score was adjusted as: sensitive (*** 12-24 positives); medium (** 6-11 positives) or poor (* 1-5 positives). Reactions with single primer sets were compared to combinations of 2 sets using M1800 at 65 (A) and 60 °C (B). Reactions with 4 different dual primer set combinations (S10+S11, S10+S12, S11+S12 and N2+E1) in M1800 (C) and E1700 (D).

**Figure 3.**
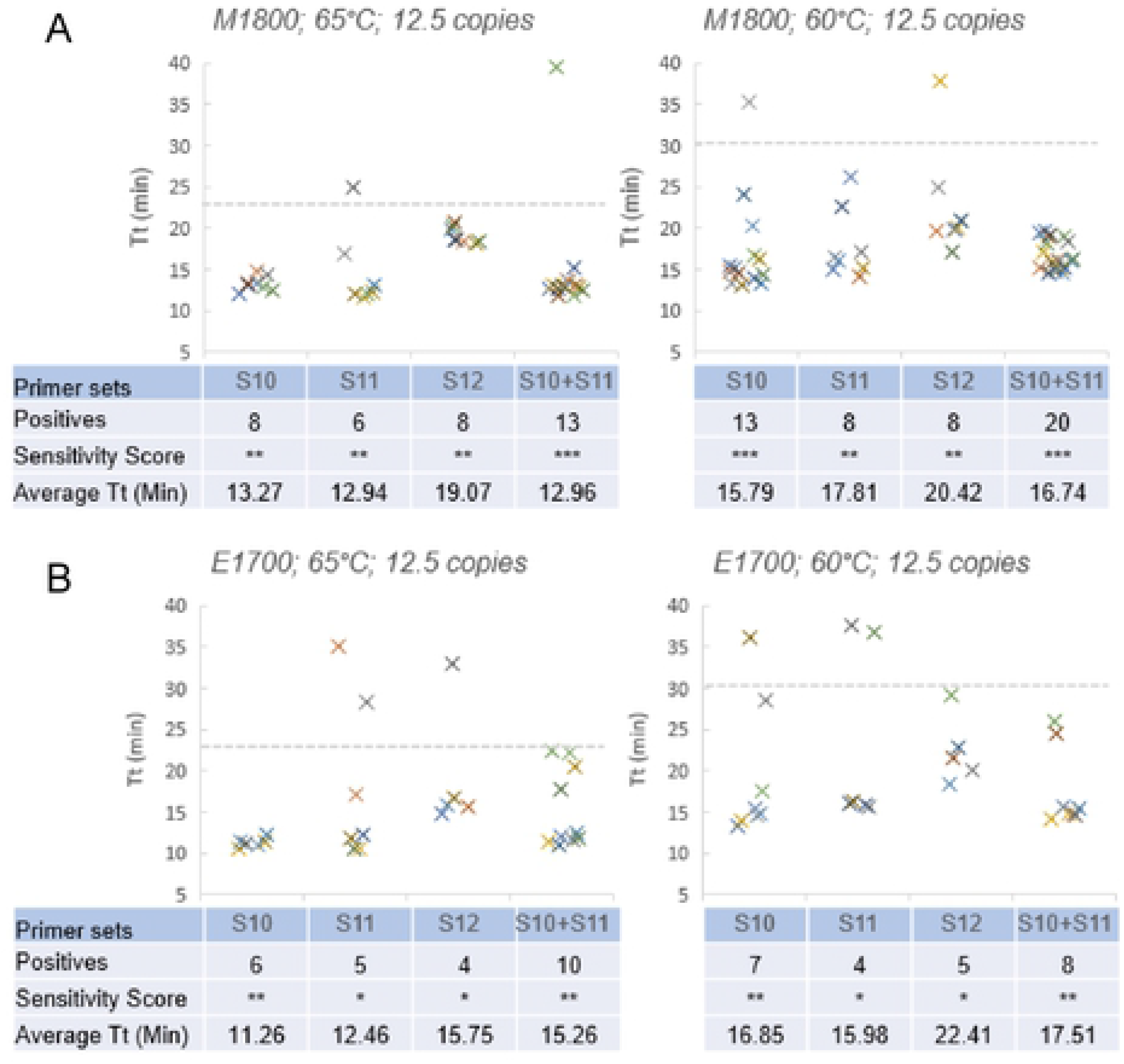
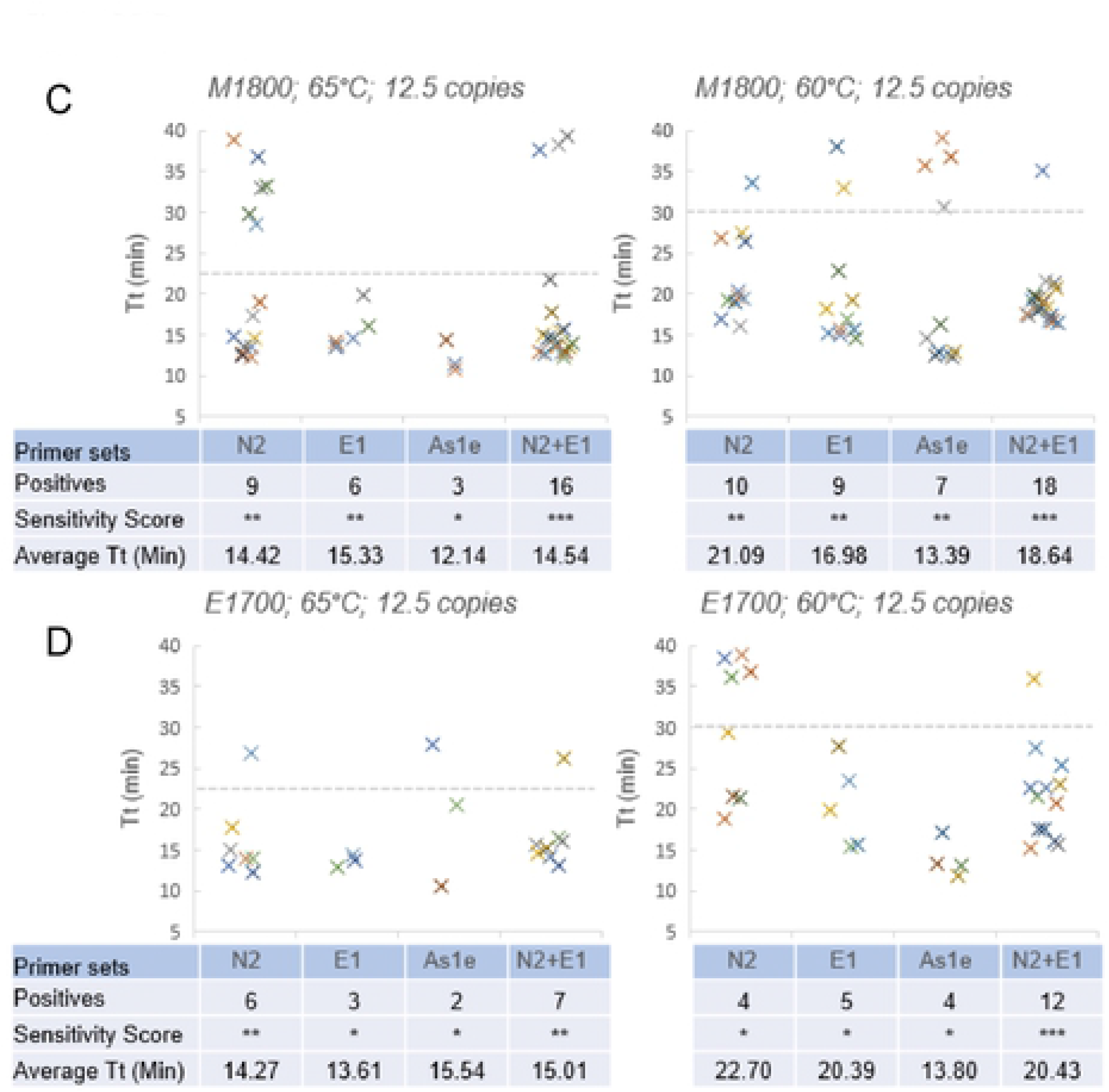
Effect of combining 2 primer sets in the presence of extremely low SARS-CoV-2 RNA template, with similar test design as Figure 2 but using only ~12.5 copies of SARS-CoV-2 RNA template. Single primer set reactions versus a combination of 2 sets using either M1800 (A, C) or E1700 (B, D) LAMP mix.

### Combination of 3 primer sets

As use of 2 primer sets showed better detection sensitivity than a single set, we evaluated whether adding a third set could further improve the detection sensitivity. We assayed two groups (S10, S11, S12; and N2, E1, As1e) with all pairwise combinations of primer sets within each group versus reactions containing all 3 sets. LAMP reactions were performed with ~12.5 copies of SARS-CoV-2 RNA template. Overall, the benefit of 3 primer sets over 2 sets was not consistent across different primer combinations. For S10+S11+S12 (Figure 4A-B), there seemed to be no advantage and it gave similar number of positives as those reactions with only 2 sets of primers combinations (S10+S11, S10+S12 and S11+S2) in both RT-LAMP reagents and at either reaction temperature. For N2+E1+As1e (Figure 4C-D), there was some advantage, and it produced more positives than those reactions containing 2 primer sets (N2+E1, N2+As1e and E1+As1e) with M1800 especially at 60 °C but not with E1700. While of course it is possible that other combinations of 3 primer sets may provide a sensitivity benefit to RT-LAMP, the results seen here indicate diminishing returns adding a 3^rd^ set to a combination of 2 primer sets, and that a combination of 2 sets is likely a sufficient approach to increasing sensitivity.

**Figure 4.**
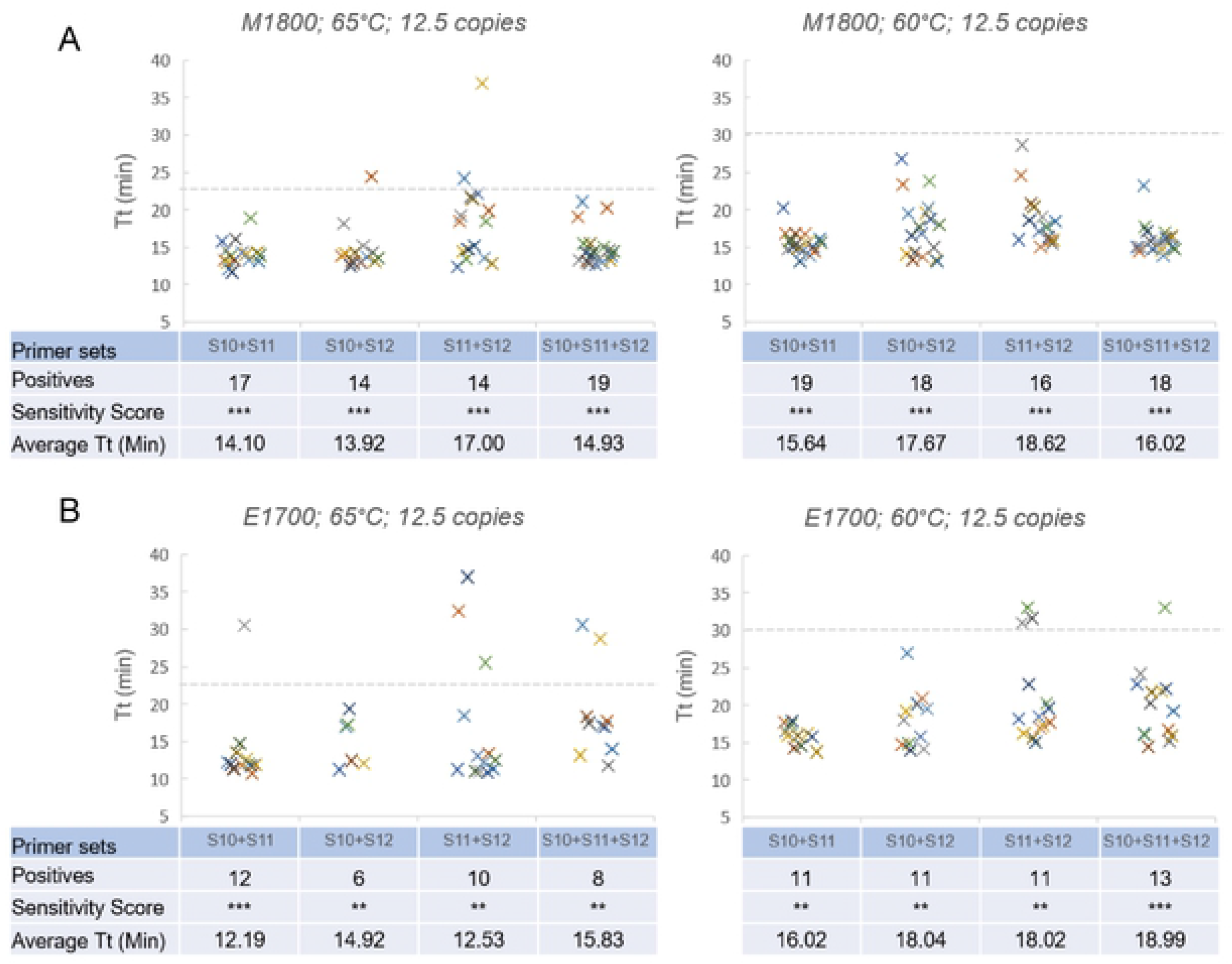
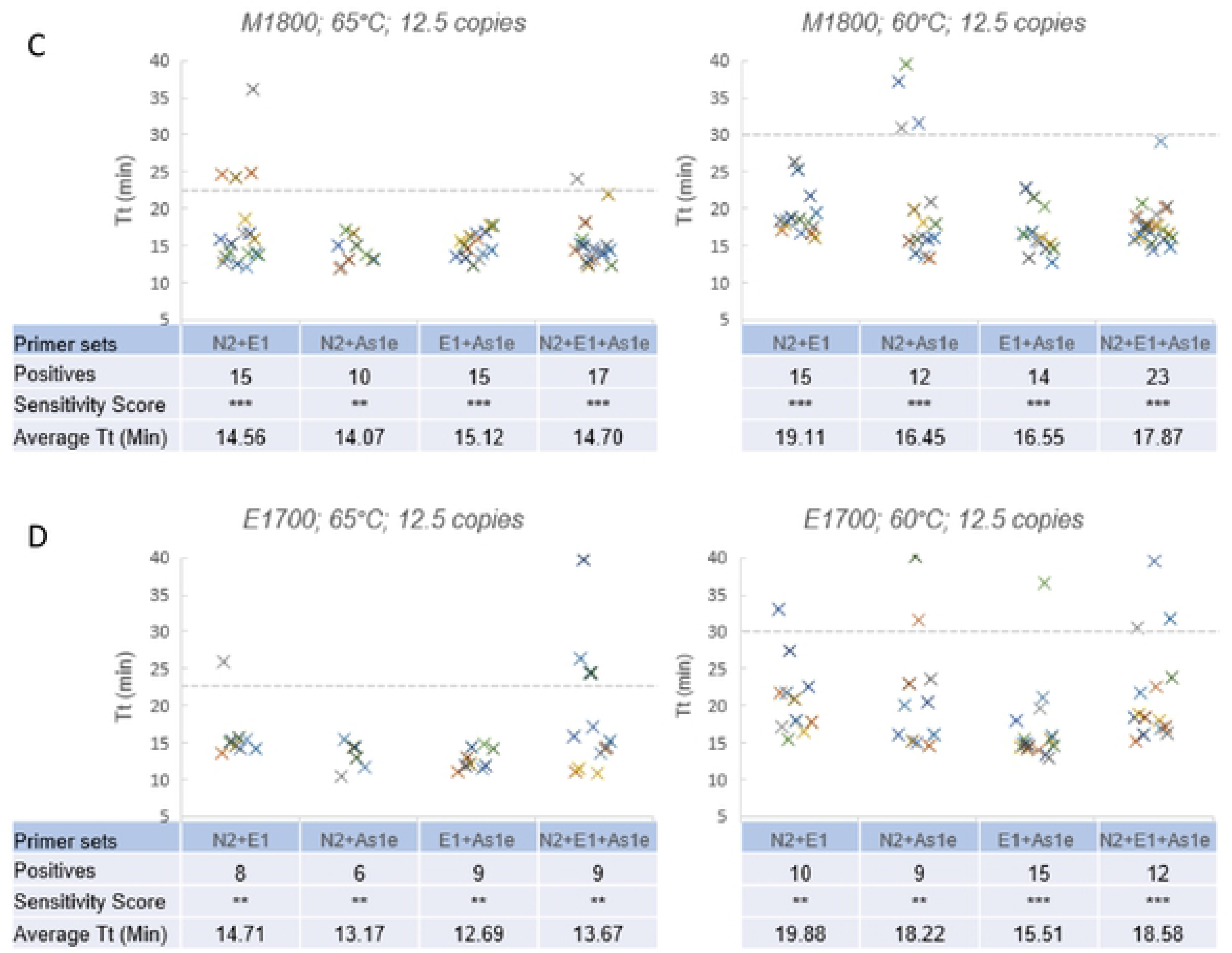
Effect of combining 3 primer sets. Detection sensitivity by a combination of 2 primer sets was compared with that by a combination of 3 primer sets in the presence of ~12.5 copies of template RNA. Combinations of 2 primer sets with versus a combination of all 3 sets using M1800 (A, C) or E1700 (B, D) LAMP mix.

## Discussion

Through the evaluation of 18 SARS-CoV-2 primer sets, we have a better understanding of the relative performance of these primers. Among them, 6 primer sets (S10, S11, S12, S13, N2 and E1) gave the most sensitive SARS-CoV-2 RNA detection with two LAMP formulations and temperatures. 2 primer sets (S4 and S18) showed high sensitivity with M1800 Colorimetric LAMP only at 65 °C, and 2 sets (S17, Joung) showed high sensitivity at 60 °C with both M1800 and E1700. Some of these primer sets were also shown to be among the most sensitive in previous comparisons. For example, S4, S10, S11, S13 were among the 6 final sets in Dong et al (along with S14 and S17) [8]. Their study gave more weight of scoring to primer sets that had faster reaction speed instead of percentage of positives, but we observed in comparison here minimal differences in speed across the sets and little correlation with speed and sensitivity. S11 and S18 appeared in the final 5 along with N-Baek, S-Huang and As1e in Janikova et al [9] though this work assayed only a small number of reactions in comparing the sets.

Overall we did observe some consistent trends, with a slight improvement in sensitivity with the M1800 Colorimetric LAMP as compared to the general-purpose E1700. Optimal reaction temperature varied for the 18 sets. Using M1800 Colorimetric LAMP, 13 sets showed similar performance at the two temperatures, 3 more sensitive at 60 °C and 2 more sensitive at 65 °C. Using E1700 there was more variation, with 6 sets giving similar results, 7 more sensitive at 60 °C and 5 more sensitive at 65 °C.

We further showed that the detection sensitivity could be increased by combining 2 sensitive primer sets into the same RT-LAMP reaction, for both temperatures and LAMP formulations evaluated. This strategy was described in our previous study and the increase is likely a result of combined probability of detection in the reaction [10]. All pairwise combinations of 2 primer sets showed increased detection sensitivity than any single primer set. A comparison of 4 sets of 2 primer combinations (S10+S11, S10+S12, S11+S12 and N2+E1) showed similar detection sensitivity in both M1800 and E1700 at both temperatures. We also tested combining 3 sets of primers into the same reaction. One group (N2+E1+As1e) seemed to give further increased detection sensitivity as we found before [10] but not with the other group (S10+S11+S12). The use of 2 sets together consistently enhanced sensitivity and could be considered for any application where that is important, with the use of 3 primer sets a possible additional step but less likely to provide a benefit. In addition to increasing sensitivity, combining primer sets for different gene targets reduces the potential impact of sequence mutations and variants that may arise in the targeted areas. While the study presented here is limited to synthetic RNA control templates, the choice of optimal primer set and the sensitivity benefit of primer set combination is a fundamental starting point for any diagnostic assay. SARS-CoV-2 unfortunately remains a significant public health concern with continuing needs for diagnostic and surveillance testing, increasingly in field, point-of-care, and even home settings removed for traditional clinical laboratory infrastructure where RT-LAMP is particularly valuable. Coupled with proper validation of different sample sources and processing methods, these conditions and recommendations could significantly improve the diagnostic sensitivity of detecting SARS-CoV-2 and any future diagnostic targets with RT-LAMP.

